# SMRT sequencing reveals differential patterns of methylation in two O111:H- Shiga toxigenic *Escherichia coli* isolates from a historic hemolytic uremic syndrome outbreak in Australia

**DOI:** 10.1101/173336

**Authors:** Brian M. Forde, Lauren J. McAllister, James C. Paton, Adrienne W. Paton, Scott A. Beatson

**Affiliations:** Australian Infectious Diseases Centre, The University of Queensland, Brisbane, Australia; The School of Chemistry and Molecular Biosciences, The University of Queensland, Brisbane, Australia.; Research Centre for Infectious Diseases, Department of Molecular and Cellular Biology, University of Adelaide, Adelaide, SA 5005, Australia.; Australian Centre of Ecogenomics, The School of Chemistry and Molecular Biosciences, The University of Queensland, Australia.

## Abstract

Shiga toxigenic *Escherichia coli* (STEC) are important food-borne pathogens and a major cause of haemorrhagic colitis and haemolytic-uremic syndrome (HUS) worldwide. In 1995 a severe HUS outbreak in Adelaide occurred. A recent genomic analysis of STEC O111:H-strains 95JB1 and 95NR1 from this outbreak found that the more virulent isolate, 95NR1, harboured two additional copies of the Shiga toxin 2 (Stx2) genes although the structure of the Stx2-converting prophages could not be fully resolved due to the fragmented assembly. In this study we have used Pacific Biosciences (PacBio) single molecule real-time (SMRT) long read sequencing to characterise the complete epigenome (genome and methylome) of 95JB1 and 95NR1. Using long reads we completely resolved the structure of two, tandemly inserted, stx2-converting phage in 95NR1. Our analysis of the methylome of 95NR1 and 95JB1 identified hemi-methylation of a novel motif (5’-C<u>T</u>GC^m6^AG-3’) in more than 4000 sites in the 95NR1 genome. These sites were entirely unmethalyted in the 95JB1, including at least 180 potential promoter regions that could explain regulatory differences between the strains. We identified a Type IIG methyltransferase encoded in both genomes in association with three additional genes in an operon-like arrangement. IS*1203* mediated disruption of this operon in 95JB1 is the likely cause of the observed differential patterns of methylation between 95NR1 and 95JB1. This study demonstrates the enormous potential of PacBio SMRT sequencing to resolve complex prophage regions and reveal the genetic and epigenetic heterogeneity within a clonal population of bacteria.

## Introduction

In 1995 a large outbreak of hemolytic uremic syndrome (HUS) occurred in Adelaide, South Australia. This outbreak was attributed to uncooked, fermented, dry sausage contaminated with Shiga toxigenic *Escherichia coli* (STEC) O111:H-(Paton et al. 1996). A total of 23 cases were confirmed, all in children aged 6 months - 14 years. As a result of infection sixteen of the 23 children required dialysis and one 4 year old child died. STEC isolates from the outbreak were found to be highly virulent with an infectious dose requiring as little as 1 organism per 10g of sausage. Interestingly, strains isolated after January 25^th^ 1995 appeared more virulent and, based on Southern blot analysis, were predicted to encode a second copy of *stx*2AB (Centers for Disease and Prevention 1995; Paton et al. 1996).

Recently, the 1995 Adelaide outbreak was re-examined using genome sequencing (McAllister et al. 2016). Two strains, 95JB1 and 95NR1 isolated pre and post January 25th 1995, respectively, were sequenced on the Illumina GAII platform and assembled *de novo*. Comparison of the draft genomes of 95JB1 and 95NR1 identified a Stx1-converting prophage and Stx2-converting phage shared by both strains as well as ~50 kb of phage-associated sequence that was present in 95NR1 but absent in 95JB1. Based on read coverage and long-range PCR it was inferred that there were two additional Stx2 prophage in 95NR1 when compared to 95JB1 (McAllister et al. 2016). All single nucleotide polymorphisms (SNPs) differentiating 95JB1 from 95NR1 had occurred in 95JB1 after divergence suggesting that the two additional Stx2 prophages were likely deleted from 95JB1 rather than acquired by 95NR1 (McAllister et al. 2016). However, due to the inability of short read sequencing technologies to accurately resolve repetitive loci, the structure and genomic context of these prophages could not be unambiguously resolved.

Due to the complexity of repeat sequences such as insertion sequences (IS), bacterial genome assemblies generated from short sequencing reads are usually highly fragmented (Koren et al. 2013). Resolving these fragmented assemblies requires time consuming and costly finishing procedures which are beyond the scope of most research projects (MacLean et al. 2009; Nagarajan et al. 2010). Consequently, these genomes are often submitted as unfinished drafts with many containing misassemblies and platform-specific sequencing errors (Phillippy et al. 2008; Chain et al. 2009). Fragmented assemblies are unsuitable for studying structural variation and important information relating to the copy number and context of IS elements and phage can be difficult to discern. The Pacific Biosciences (PacBio) Single Molecule Real Time (SMRT) sequencing platform is able to completely resolve most bacterial genomes by producing reads of sufficient length to span complex repeat loci and generate complete assemblies without the need for costly manual finishing (Eid et al. 2009) (Chin et al. 2013; Koren et al. 2013; Brown et al. 2014; Forde et al. 2014).

A remarkable feature of SMRT sequencing is the capacity to determine the methylation status of every sequenced nucleotide (Clark et al. 2012). DNA methylation is the most common post replicative modification in bacteria (Roberts et al. 2003) and is known to influence a wide variety of host processes, including DNA replication, repair and transcriptional regulation (Blow et al. 2016). Until recently the lack of a simple, efficient method to determine the methylation status of DNA has resulted in these modifications being largely ignored in the bacteria. However, by monitoring the actions of single DNA polymerase enzymes in real-time SMRT sequencing can simultaneously determine nucleotide sequence and the methylations status of each nucleotide.

Methylation is mediated in bacteria by methyltransferase (MTase) enzymes. MTase are found as components of restriction-modifications (RM) systems, which in their simplest form are generally composed of a MTase and a restriction enzyme (REase), although some classes of MTase are able to catalyse both restriction and methylation (Roberts et al. 2003). RM systems are ubiquitous in bacteria and highly diverse, but until recently a small number have been characterised. The primary role of RM systems is to protect the host genome from invasion by foreign DNA, such as transposons, phage and plasmids (Takahashi et al. 2002). MTase-genes can also occur independently of RM systems. These ‘orphan’ MTase are common in bacterial genomes and can be involved in genome regulation (Blow et al. 2016). The most well characterised orphan MTase is Dam (Casadesus and Low 2006). Dam is an established regulator of gene expression in *E. coli* and also has a role in pathogenicity (Boye and Lobner-Olesen 1990; Low et al. 2001; Casadesus and Low 2006). Less well studied MTases that are components of RM systems may also have roles beyond genome defence. For example, Type III (Res, Mod) RM systems are known to be involved in phase variation in a number of bacterial species (de Vries et al. 2002; Fox et al. 2007; Seib et al. 2015) and phase variation of Type IIG RM systems has been shown to alter gene expression in *Campylobacter jejuni* (Anjum et al. 2016).

To investigate the differences between two STEC O111:H-outbreak strains 95JB1 and 95NR1 we determined their complete genome assemblies and methylomes using PacBio SMRT sequencing. We completely resolve the genetic structure of all prophage-encoding regions, including two distinct, but closely related, Stx2-converting prophage in 95NR1 that are tandemly inserted in the same genomic location and appear to have been deleted from 95JB1. We identify all putative MTases in both strains, define their target sequences and show the activity of a previously uncharacterised methyltransferase in 95NR1 that is not active in 95JB1. Furthermore, we unambiguously determine the copy number and context of all IS elements in the genomes of 95NR1 and 95JB1 and reveal that a single difference in the IS complement could be directly responsible for differences in their methylome profiles.

## Results

### Complete genome assembly reveals full sequence of tandemly arrayed Stx2 prophages in 95NR1

*De novo* assembly of *E. coli* O111:H-strains 95JB1 and 95NR1 generated single circular chromosomal contigs of 5,347,879 bp and 5,467,946 bp, respectively. Previously reported short read assemblies of 95NR1 (accession: AVDU00000000.1) and 95JB1 (accession: AWFJ00000000.1) contained 182 and 179 contigs, respectively, with contig N50 sizes of less than 100 kb (McAllister et al. 2016) (Table 1). A SNP comparison of the complete genomes of 95JB1 and 95NR1 is in agreement with previous observations using Illumina data (McAllister et al. 2016). Using *E. coli* O111:H-strain 11128 as a reference, McAllister *et al* identified six SNPs, 5 on the chromosome and one plasmid encoded SNP, which discriminate 95JB1 from 95NR1 (McAllister et al. 2016). Our SNP comparison of the complete PacBio assemblies of 95JB1 and 95NR1 identified four of the five chromosomally located 95JB1 SNPs and the single plasmid encoded SNP (Table 2). The remaining chromosomal SNP was located in a phage tail protein and due to the repetitiveness of these proteins it was not considered reliable for strain discrimination.

**Table 2.** SNPs differentiating 95NR1 and 95JB1.

| Strain | Base Change | 95JB1 site | 95NR1 site | Amino Acid Change | Illumina* | PacBio | Annotation |
| --- | --- | --- | --- | --- | --- | --- | --- |
| 95JB1 | C-T | 587038 | 587038 | P-L | + | + | *glxK* (Glycerate kinase II) |
| 95JB1 | G-T | 3227026 | 3227373 | V-F | + | + | End of Stx1 prophage |
| 95JB1 | G-A | 3602596 | 3602945 | E-K | + | + | *metK* (Methionine adenosyltransferase 1) |
| 95JB1 | G-C | 3813994 | 3814343 | Stop codon-Y | + | + | *fadH* (2,4-dienoyl-CoA reductase) |
| p95JB1B | G-C | 20482 | 20477 | P-A | + | + |  |

\* SNPs identified by McAllister et al (McAllister et al. 2016)

**Table 1:** Assembly Statistics.

| Strain | Number of SMRTCells | Number of PacBio contigs | PacBio contig N50 (bp) | Number of Illumina Contigs | Illumina contig N50 (bp) |
| --- | --- | --- | --- | --- | --- |
| 95JB1 | 2 | 4 | 5373164 | 179 | 91606 |
| 95NR1 | 2 | 3 | 5462770 | 182 | 94405 |

The ~120 kb difference in chromosome size between 95JB1 and 95NR1 was largely attributed to two tandemly inserted Stx2-converting prophages in 95NR1 (Phi14 and Phi15) that are absent from 95JB1 (Figure 1a). Phi14 and Phi15 from 95NR1 are highly conserved and share 99% nucleotide sequence identity across 79% of their genomic sequence with regions of difference largely confined to the 5’ end of the prophage sequences (Figure 1b). This pattern of sequence identity explains why assembly of this region was not possible using short read data alone. Indeed, the structure and context of all phage-encoding regions, including the Stx2 and Stx1 converting prophage (Phi10 and Phi11, respectively) carried by both outbreak strains, (Table S1) were readily resolved in the complete PacBio assemblies.

**Figure 1:**
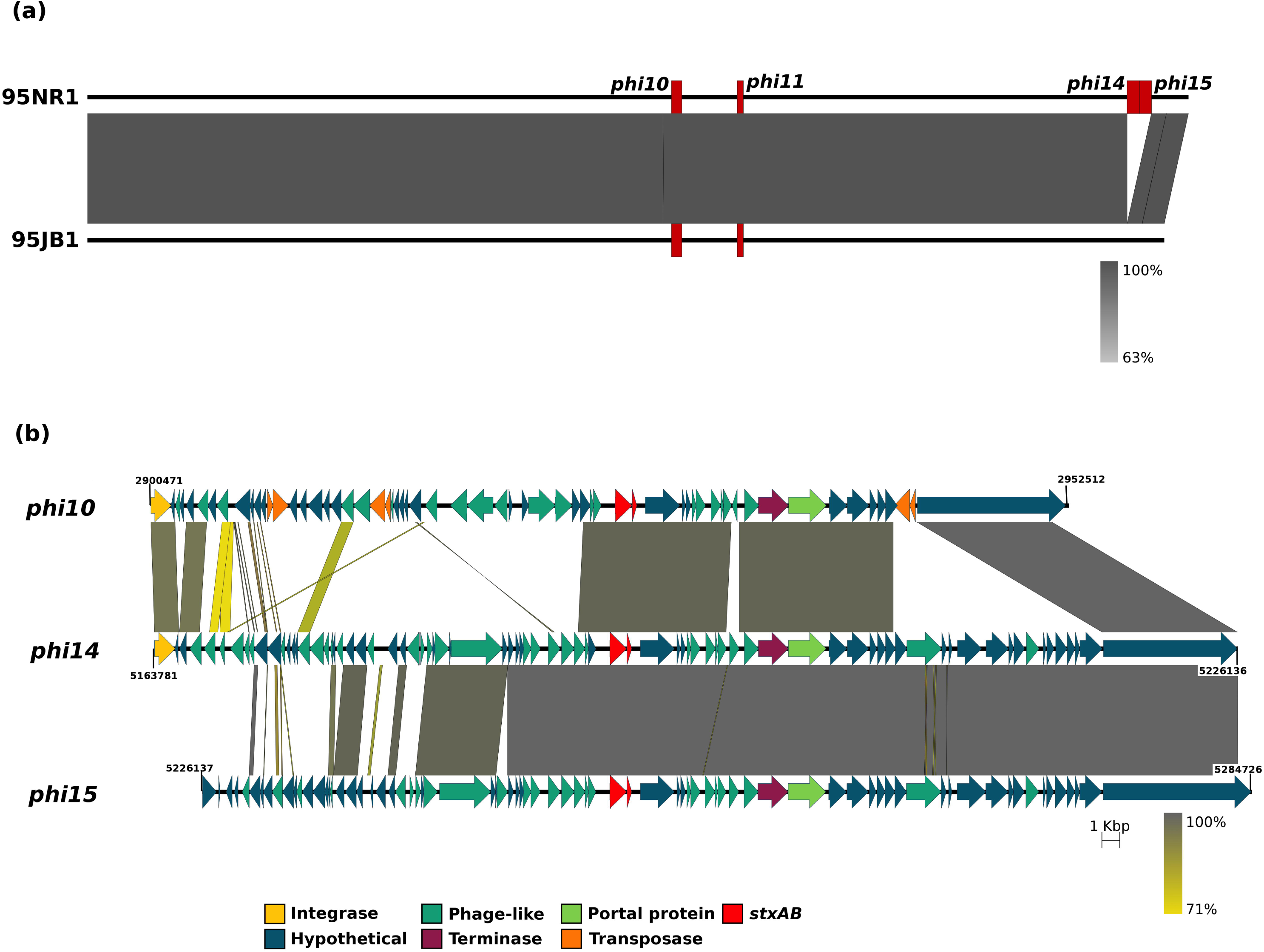
Comparison of *E. coli* 95JB1 and 95NR1 highlighting the position and context of Stx-carrying prophage. (a) Pairwise nucleotide comparison of 95NR1 (top) and 95JB1 (bottom) chromosomes. The chromosomes of 95NR1 and 95JB1 are represented to scale by the black bar with the Stx-carrying prophage insertion points indicated with red rectangles. Popouts display schematic representations of the four Stx-carrying phage carried on one or both strains. (b) Pairwise nucleotide comparison of three Stx2-converting phage from 95NR1. Phi10, Phi14 and Phi15 are represented to scale. Prophage encoded proteins are represented by arrows coloured as per the legend. Yellow and grey shading between phage represent regions of homology from 71% (yellow) to 100% (grey) nucleotide sequence identity. Prophage genes are coloured as per the legend.

Two plasmids, similar to the P1 and EHEC plasmids from *E. coli* 11128 (Ogura et al. 2009), were completely assembled in the PacBio assemblies of 95JB1 and 95NR1 (p95NR1A/p95JB1A and p95NR1B/p95JB1B, respectively). In contrast to the draft Illumina assemblies of 95JB1 and 95NR1 (McAllister et al. 2016), the small pO111_4 and pO111_5 colicin plasmids were not detected in the PacBio assemblies or in the raw PacBio read data consistent with their exclusion during the library preparation process.

### Insertion Sequence profiles differ between outbreak strains

To explore if there were any additional mobile genetic element differences between the outbreak strains we first compared the complete plasmids of each strain. Whereas the P1 plasmids were identical in both strains, the EHEC plasmid, p95JB1B, contained an additional IS*3* ssgr IS*51*-family insertion sequence (99% nucleotide sequence identity to IS*1203* from *E. coli* O111:H-PH) that was not present in p95NR1B (data not shown). This IS*1203*-like element has inserted into the 3’ end of the transposase EC95JB1_B00047 within an IS*91*-like insertion element.

We next surveyed the chromosomal Insertion Sequence (IS) profiles of each outbreak strain (Figure 2). Both 95JB1 and 95NR1 contain 17 different families of IS elements with two or more copies on their respective chromosomes (Table S2). Notably, 95JB1 encodes an additional chromosomal copy of an IS*3* ssgr IS*51*-family element (100% identical to the additional IS*1203*-like element on p95JB1B) inserted at the 3’ end of EC95JB1_03899 (Table S3). EC95JB1_03899 is predicted to encode an ATPase. However, the IS has inserted ~150 bp upstream of the translational start site of a putative methyltransferase (MTase) gene (EC95JB1_03895), suggesting a possible functional role due to a polar effect on transcription.

**Figure 2:**
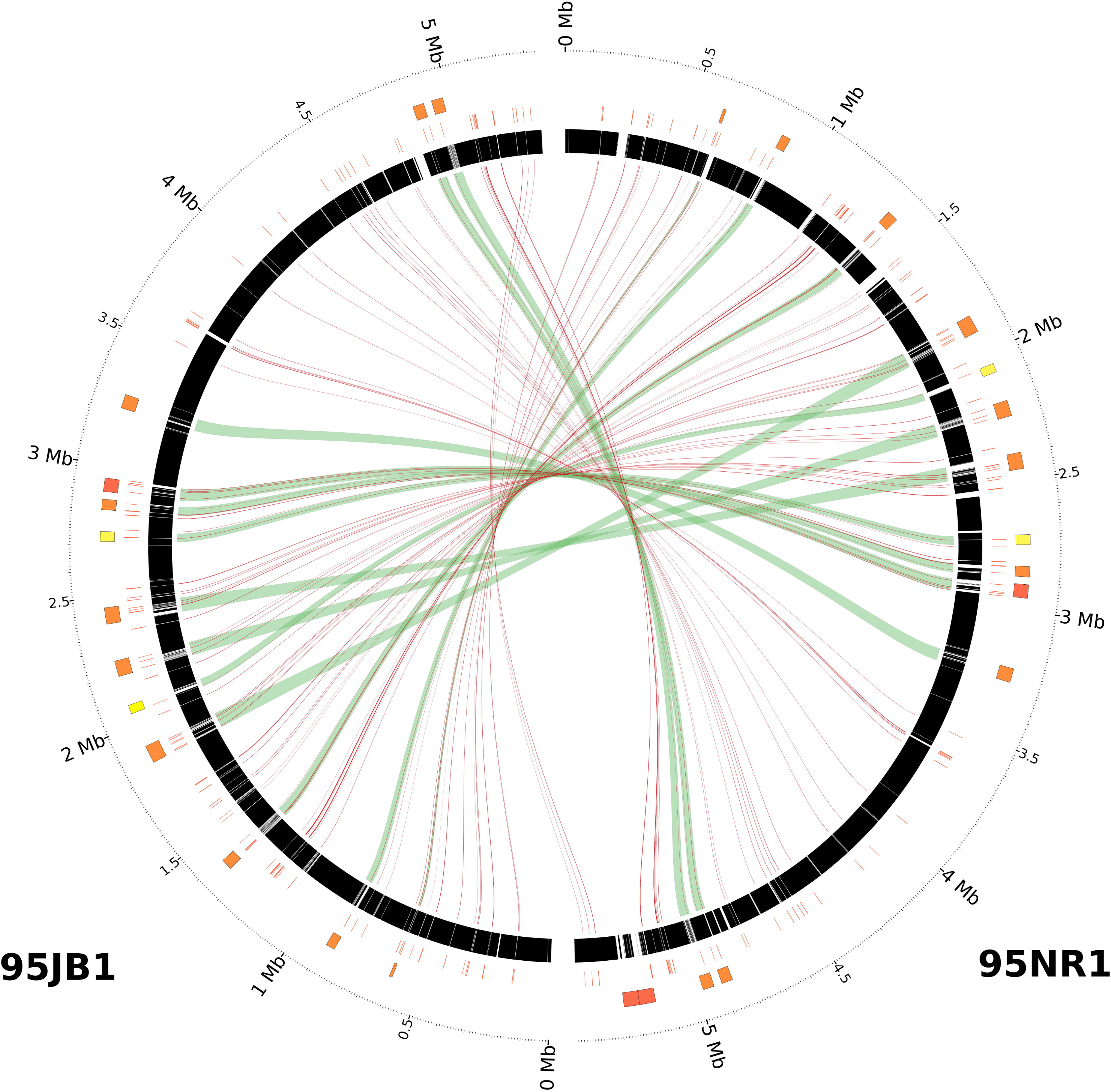
Comparison of 95JB1 and 95NR1 genome assemblies. Circos plot comparing prophage and IS content of 95JB1 (left) and 95NR1 (right). Putative prophage regions are highlighted on the outer most track by coloured rectangles: Stx-converting (Red), myoviriadae (yellow), other (orange). Small, degenerate phage–associated regions are highlighted on the outer most track in grey. The position and context of all IS elements are represented by red bars on the middle track. Draft Illumina assemblies of 95JB1 and 95NR1, mapped to their complete genomes, are represented in black on the inner most ring where assembly gaps are shown as white space. Green ribbons connect prophage sequences common to both strains. Red lines connect IS that are common to both strains.

### Three additional MTases are encoded on the additional 95NR1 Stx2 prophages

Bioinformatic characterisation of 95JB1 revealed that the strain encodes seven putative MTases in addition to Dam (target site: 5’-G^m6^ATC-3’) and Dcm (target site: 5’-C^m5^CWGG-3’). Five of these six putative MTases correspond to enzymes with known specificity that have been previously characterised in the O104:H4 serotype *E. coli* outbreak strain C227-ll (Fang et al. 2012). These include homologs of the C227-11 orphan MTases M.EcoGI, M.EcoGII, M.EcoGVI and M.EcoGIX (M.Eco95JB1I, M.Eco95JB1II, M.Eco95JB1V and M.Eco95JB1VII, respectively), the Stx phage-borne M.EcoGIII (M.Eco95JB1III), and the Dam paralog M.EcoGV (M.Eco95JB1IV); the remaining MTase, M.Eco95JB1VI, is homologous to Type IIG RM system SenTFIV from *Salmonella enterica* subsp. enterica serovar Typhimurium (76% amino acid similarity)(Table 3).

**Table 3.**
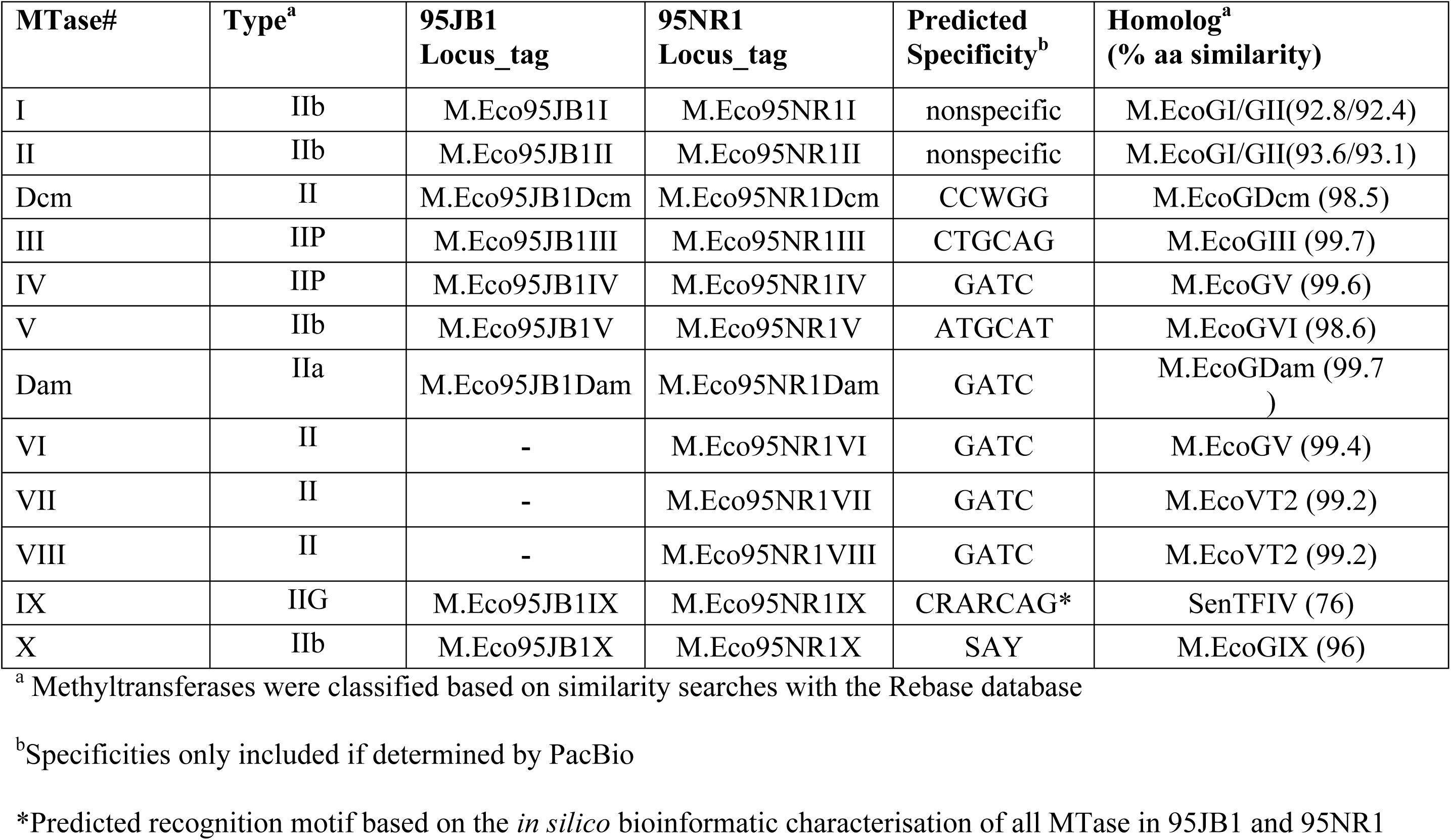
Summary of Methyltransferase genes from the chromosomes of 95JB1 and 95NR1.

In addition to encoding all MTase-encoding genes identified in 95JB1, 95NR1 also encodes three additional Dam homologs encoded on its additional Stx-converting prophages (Table 3). These comprise: (i) an additional copy of the Dam homolog M.EcoGV located on Stx2-converting prophage Phi14 (M.Eco95NR1VI), (ii) an identical copy of the orphan MTase M.EcoVT2Dam of *E. coli* prophage VT2-Sa on Stx2 prophage Phi14 (M.Eco95NR1VII) and (iii) a second identical copy of M.EcoVT2Dam on Stx2 prophage Phi15 (M.Eco95NR1VIII).

### The methylome of 95JB1 and 95NR1 show remarkably different methylation patterns

Using the in-built capacity of PacBio to detect methylated nucleotides we detected two distinct recognition motifs (5’-G^m6^A<u>T</u>C-3’ and 5’-C<u>T</u>GC^m6^AG-3’) in 95JB1 and 95NR1, that match MTases with known specificities. 5’-GATC-3’ is a well characterised methylation motif, routinely identified in *E. coli* methylome analyses (Fang et al. 2012; Powers et al. 2013; Cooper et al. 2014; Forde et al. 2015) and known to be targeted by Dam (Geier and Modrich 1979). We predict that the 5’-CTGCAG-3’ motif is targeted by EC95JB1_EC95NR1_MTases in 95JB1 and 95NR1 (M.Eco95JB1III/M.Eco95NR1III) that share 99% amino acid identity with M.EcoGIII, a previously characterised *Pst*I homolog shown to methylate 5’-CTGCAG-3’ (Fang et al. 2012). No additional methylated motifs in 95JB1 were detected suggesting that the remaining putative MTases are inactive under the tested conditions (Table 4). Dcm-mediated methylation of cytosine (5’-C^m5^CWGG-3’) could not be detected using the method used in this study.

**Table 4.** MTase recognition motifs identified in 95JB1 and 95NR1.

| **Motif** | **Modification Type** | **Number Detected** | **Number in Chromosome** | **Methylated (%)** | **Mean IPD Ratio** |
| --- | --- | --- | --- | --- | --- |
| **95JB1** |  |  |  |  |  |
| CTGCAG | m6A | 2390 | 2390 | 100.0 | 6.9934<br>053 |
| GATC | m6A | 41658 | 41686 | 99.9 | 5.6757<br>846 |
| **95NR1** |  |  |  |  |  |
| CTGCAG | m6A | 2434 | 2434 | 100.0 | 7.1674<br>423 |
| CRARCAG | m6A | 4074 | 4074 | 100.0 | 7.1986<br>117 |
| GATC | m6A | 42242 | 42270 | 99.9 | 5.7941<br>27 |

Remarkably, 95NR1 also contained more than 4000 methylated bases corresponding to a third motif, 5’-CRARC^m6^AG-3’ (Table 4). As no methylation was detected at the corresponding motifs in 95JB1 we considered that this new activity must relate to an MTase that is not present or non-functional in 95JB1. Screening the 5’-CRARCAG-3’ motif against REBASE (Roberts et al. 2010) confirmed that a cognate MTase enzyme has not been previously characterised and to the best of our knowledge it represents a novel MTase target recognition site (Table 4).

### A novel MTase responsible for methylation of the CRARCAG motif

In order to identify the MTase responsible for methylation of the 5’-CRARCAG-3’ motif in 95NR1 (but not 95JB1) we first examined the specificities of the experimentally determined *E. coli* C227-11 MTases for which there are five close homologs in 95NR1/95JB1 (>92% amino acid sequence identity; Table 3) (Fang et al. 2012). Based on the high amino acid identity between these homologs it is highly unlikely that any C227-11 MTase homologs could be responsible for methylation of the 5’-CRARCAG-3’ motif. Similarly, M.EcoVT2Dam is known to target 5’-GATC-3’; thus both homologs of M.EcoVT2Dam in 95NR1 (99% amino acid identity) would also be expected to target 5’-GATC-3’ (Radlinska and Bujnicki 2001). The remaining candidate MTase encoded by M.Eco95NR1IX shares 76% amino acid identity to SenTFIV, a Type IIG R-M system which targets the GATCAG recognition site. Minor difference in the target recognition domain (TRD) of homologous MTase enzymes can result in major differences in target specificity as exemplified in *E. coli* EC958 where the Type IIG R-M system M.EcoMVII and SenTFIV share 68% amino acid identity, but their target sites are very different (5’-CANCATC-3’ and 5’-GATCAG-3’, respectively). Furthermore, 100% hemi-methylation of the 5’-CRARCAG-3’ motif as observed in 95NR1 is characteristic of the Type IIG family of MTases. Based on these observations, we propose that the MTase M.EcoEC95NR1_IX catalyses CRARCAG modification in 95NR1 and that the lack of corresponding methylation in 95JB1 is due to IS-mediated transcriptional inactivation of the orthologous M.Eco95JB1VI gene.

### IS insertion results in inactivation of M.Eco95JB1IX

BLAST comparison (blastp, E-value cutoff=1e^−3^) of M.Eco95NR1IX against a database of complete *E. coli* genomes identified homologs of this MTase in four additional *E. coli* strains: *E. coli* str. HS (EcHS_A0339; 76% amino acid sequence identity), *E. coli* O111:H-str. 11128 (ECO111_5156; 100% amino acid sequence identity), *E. coli* str. ‘clone i2’ (i02_4877; 76% amino acid sequence identity) and *E. coli* str. ‘clone i14’ (i14_4877; 76% amino acid sequence identity). With the exception of ECO111_5156, homologs of M.Eco95NR1IX exhibited extensive variation in their TRDs and are likely to have different target specificities (Figure S1a). Interestingly genes upstream of M.Eco95NR1IX are highly conserved in all strains (Figure S1b) and suggest that M.Eco95NR1IX could be transcribed as part of a large 8.5 kb operon encoding a Type IIG MTase (M.Eco95NR1IX), a putative ATPase (EC95NR1_04072) and two hypothetical proteins whose function is currently unknown (EC95NR1_04073 and EC95NR1_04074) (Figure S2; Table S4). A putative promoter region (5’-TGGCAT-14 bp-CATTACAAT-3’) was identified 50 bp upstream of EC95NR1_04074 and could indicate the primary transcriptional start site for this operon. Predicted promoter regions were also identified 68 bp upstream of EC95NR1_04072 and overlapping the predicted start codon of M.Eco95NR1IX. An additional predicted promoter region was observed in 95JB1 which was located on the boundary of the IS insertion into EC95JB1_03895 (Figure S2). Based on these observations we propose that the IS insertion into EC95JB1_03899 results in premature transcriptional termination of the operon leaving M.Eco95JB1IX untranscribed.

### The functional characterisation of CRARCAG methylation

To determine if methylation of 5’-CRARCAG-3’ motifs could have a functional role in the genome of 95NR1 we analysed the distribution of sites located within 300 bp upstream of an annotated start codon. We identified 871 candidate genes where 5’-CRARCAG-3’ methylation might have a regulatory role (Table S5). These include the flagellar genes *fliCDSHIJKQPR*, *flgCJG* and the flagellar regulator *flk*; Type II secretion associated genes *epsMLJHGE*; the pilus associated genes *papD* and *pilN*, Shiga toxin gene *stx*_*1*_*B* (Phi11); the two component regulatory system BaeSR, and EutR, a transcriptional regulator associated with EHEC pathogenesis (Luzader et al. 2013). Upstream sequences were further analysed for the presence of putative promoter regions using Neural Network Promoter Prediction. A total of 601 of the 871 candidate genes were found to contain putative promoter regions within 300 bp of their start codon, 177 of which contained a methylated 5’-CRARCAG-3’ motif (Table S5). Clustering of these genes based on their functional class revealed no significant enrichment in any functional category when compared to the functional clustering of all genes (Figure S3).

### Distribution of Methylated motifs reveals difference in methylation patterns within prophage sequences

There was an observed difference in the distribution of the 5’-GATC-3’ motif between the core genome and the prophages in both 95NR1 and 95JB1 (Figure 3). This difference is attributable to the presence of GATC-free regions within the prophages and is suggestive of selection against Dam methylation in certain phage genes. Differences in the distribution of GATC sites between the core and accessory genome have previously be described in *E. coli* K12 (Henaut et al. 1996) and more recently in *E. coli* EC958 (Forde et al. 2015). Interestingly, these prophage-associated GATC-free regions are enriched for the 5’-CTGCAG-3’ motif in most prophages in 95JB1 and 95NR1 with the exception of Phi9 (unclassified Myoviridae) and Phi12 (Mu-like Myoviridae) where the 5’-CTGCAG-3’ motif is significantly under-represented compared to the core genome (P <= 0.0001). In 95NR1 the 5’-CRARCAG-3’ motif appears to be randomly distributed throughout the genome and exhibited no significant enrichment bias to either the core or accessory genome.

**Figure 3:**
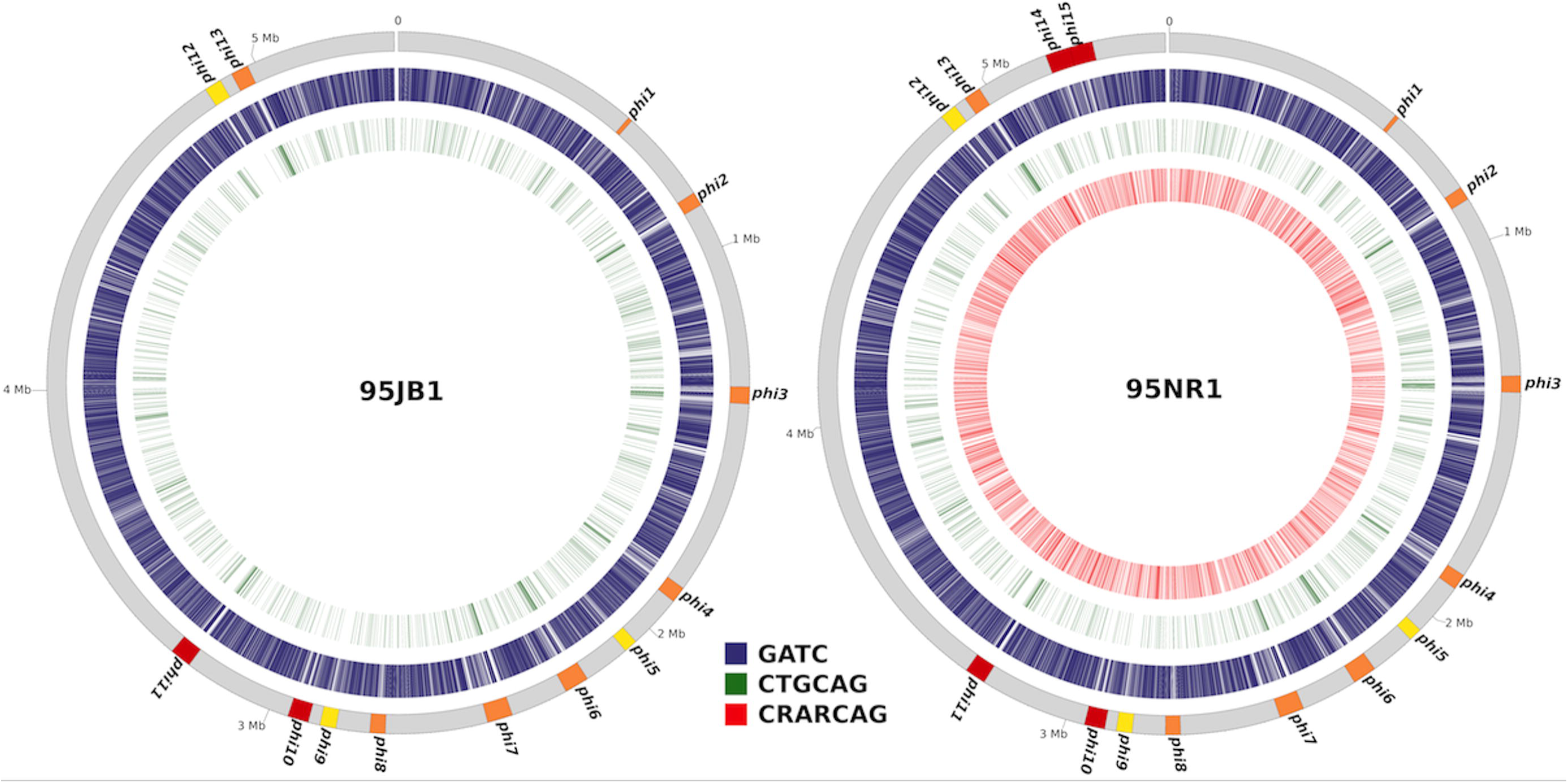
Comparison of methylated DNA on the chromosomes of 95JB1 and 95NR1. Circos plot displaying the distribution of methylated nucleotides on the chromosomes of *E. coli* 95JB1 (left) and 95NR1 (right). Prophage insertion points are highlighted on the outer-most track in orange with Stx-carrying prophage in red and myoviriadae in yellow. The remaining coloured tracks highlight the chromosomal positions of all methylated sites for each motif. Tracks are coloured as per the legend; GATC, red; CTGCAG, green; CRARCAG, red.

## Discussion

*E. coli* 95JB1 and 95NR1 are two isolates from a historical HUS outbreak in Adelaide, Australia in 1995 (Centers for Disease and Prevention 1995; Paton et al. 1996). 95NR1, the more virulent of the two strains, is characterised by the presence of two additional Stx2 prophages and two additional copies of *stx*_*2*_*AB* when compared to 95JB1 (McAllister et al. 2016). Previous analysis of the Illumina draft genomes of these outbreak strains showed that, relative to 95NR1, 95JB1 had acquired a handful of SNPs and had lost two Stx2 prophages. Despite these findings it was not possible to resolve the sequences of most mobile genetic elements (including prophages) due to the fragmented nature of the draft assembly. In this study we hypothesised that there were other genetic differences between the strains that may account for the different virulence profiles. Using SMRT sequencing to produce complete genomes of these important historical isolates enabled the complete definition of the structure and context of all prophages (including the tandemly arranged Stx prophages in 95NR1). By analysing the kinetic variation data produced during sequencing on the SMRT platform we found a difference between the methylomes of 95JB1 and 95NR1. By identifying the position and context of all IS elements in both genomes we identified an additional chromosomal copy of an IS*1203*-like element in 95JB1 that likely prevents transcription of a Type IIG MTase encoded nearby. As we are able to exclude all other MTases encoded by 95NR1 we conclude that this IS difference accounts for the methylation patterns between the strains. It has been well established that DNA methylation can play an active role in virulence and gene regulation (Heithoff et al. 1999; Low et al. 2001; Wallecha et al. 2002; Casadesus and Low 2006). By examining methylated bases that may affect promoter regions in 95NR1, but not 95JB1, we have identified a number of potential promoters that could contribute, directly or indirectly, to the increased virulence observed in 95NR1.

Previously, a detailed analysis of the Illumina draft genome assemblies of 95JB1 and 95NR1 enabled accurate determination of prophage insertion sites, plasmid content and the multiplicity of *stx* genes (McAllister et al. 2016). As expected, repeat elements were the major cause of fragmentation in the draft genomes of 95JB1 and 95NR1, with the majority of contigs terminating at IS elements or within prophage regions. Long SMRT sequencing reads easily bridge these repeat regions generating complete, high quality assemblies with little or no manual intervention. Complete genomes are a vital resource when characterising the genetic variation which exists between strains, as the positions of IS and other mobile elements can influence host biological processes, including pathogenicity and antibiotic resistance (Conlan et al. 2014; Cameron et al. 2015; He et al. 2015; Yan et al. 2015; Zowawi et al. 2015).

Determining the context of all IS in 95JB1 and 95NR1 was crucial for characterising methylome differences between both strains. Surprisingly, we found that the most likely explanation for the observed differences in methylation between the outbreak strains was the insertion of an additional IS*1203*-like element upstream of the gene encoding M.Eco95JB1IXin 95JB1. This particular IS element was first identified in *E. coli* O111:H-strain PH (Paton and Paton 1994) and has previously been associated with insertional inactivation of genes in STEC (Kusumoto et al. 1999; Okitsu et al. 2001; Suzuki et al. 2004). Several lines of evidence support our contention that M.Eco95NR1IX/M.Eco95JB1IX catalyses the methylation of the 5’-CRARC^m6^AG-3’ motif. Other than M.Eco95NR1IX there were no candidate MTases in the 95NR1 or 95JB1 genomes that could account for the methylation of the 5’-CRARCAG-3’ motif in 95NR1 alone. M.Eco95NR1IX and SenTFIV share 76% amino acid identity consistent with a different recognition site. Furthermore, the distinctive pattern of hemi-methylation of the 5’-CRARCAG-3’ motif as observed in 95NR1 is characteristic of the Type IIG family of MTases. Formal demonstration that the 5’-CRARCAG-3’ recognition motif is methylated by M.Eco95NR1IX would require methylome profiling of a 95NR1 M.Eco95NR1IX deletion mutant.

In this study we identified two active MTases common to both 95JB1 and 95NR1: Dam and the recently characterised Stx phage-encoded RM system methyltransferase M.EcoGIII, which recognises the CTGCAG motif. M.EcoGIII is known to affect the expression of 1,951 genes in *E.coli* C227-ll, provide resistance to infection by other lambda-like phage and influence growth. It is likely that M.EcoGIII occupies a similar regulatory role in 95JB1 and 95NR1 as it does in C227-11. However, determining the functional consequences of CTGCAG methylation in 95JB1 and 95NR1 requires further analysis. Interestingly, three prophage loci in C227-11 were found to be enriched for the CTGCAG motif (Fang et al. 2012) and a similar enrichment of the CTGCAG motif was also observed in most prophages of 95JB1 and 95NR1. In contrast, two prophage regions in 95JB1 and 95NR1 (Phi12 and Phi16) contained no CTGCAG motifs, suggesting selection against the presence of these sites. In C227-11 the M.EcoGIII RM system was shown to protect against infection by other lambda-like phages, but T4 phages were resistant to restriction, which could be attributed to heavy modification of their genomes (Fang et al. 2012). 95NR1 and 95JB1 do not contain any T4 phage, although Phi12 and Phi16 are members of the Myoviridae, the class of virus to which the T4 phage belong. The lack of any CTGCAG motifs in these Phi12 and Phi16 (CTGCAG motif is expected to occur by chance every 4064 bp) suggests active selection against this motif in the Myoviridae, which would presumably render them immune to digestion by M.EcoGIII.

Bacterial MTases are known to have diverse roles which include regulation of gene expression. For example, the orphan MTase Dam has been established as regulator of gene expression in other *E. coli* species and recently the Type II RM system EcoGIII was shown to directly or indirectly affect the expression of ~1900 *E. coli* genes (Low et al. 2001; Casadesus and Low 2006). Similarly, it has been suggested that Type IIG RM systems might have a role in bacterial genomes other than protection (Loenen et al. 2014). The proximity of methylated 5’-CRARCAG-3’ sites to the predicted promotor regions of 180 genes in 95NR1 suggests that methylation of this motif could directly cause differential gene expression between 95NR1 and 95JB1. We have identified several operons and regulators that may be likely candidates for differential expression, although it should be noted that putative promoter sites were predicted *in silico* and may not reflect true promotor regions. It is also worth noting that 95NR1 and 95JB1 have been classifed as non-motile (H-) according to their original serotyping results suggesting that differential expression of flagellar loci is unlikely to result in phenotypic difference (Centers for Disease and Prevention 1995). Analysis of the transcriptome using RNA-Seq will be necessary to fully define the influence of methylation on differential regulation between these otherwise very similar strains.

DNA modifications are an established cause of phenotypic heterogeneity in both isogenic and clonal populations (Casadesus and Low 2013). Phase variation or ON/OFF switching of the pyelonephritis-associated pili (Pap) operon and antigen 43 (Ag43) are two of the most well characterised examples of methylation-mediated intercellular heterogeneity in *E. coli* (van der Woude et al. 1992; van der Woude and Henderson 2008). In these examples, differences in the methylation status of GATC motifs (mediated by Dam) in the promoter regions of *pap* and *ag43* control ON/OFF expression of these loci and intercellular heterogeneity within the clonal population. Differences in the expression of the same MTase in 95NR1 and 95JB1 (M.Eco95NR1IX and M.Eco95JB1VI, respectively) is a clear example of epigenetic heterogeneity which has arisen within this clonal population. Whether this observed epigenetic heterogeneity is driving phenotypic difference within the population is currently unknown and represents an interesting avenue for further research.

During our characterisation of the 95NR1 and 95JB1 methylomes we made the surprising discovery that a Type IIG MTase carried by 95NR1 and 95JB1 is likely encoded as part of an operon. To the best of our knowledge a Type IIG MTase has not previously been reported as a component of a multi-gene system. The function of the other proteins in this putative operon and whether they are linked to the activity of the MTase is currently not known and requires further research which is beyond the scope of this study. Despite identifying a putative primary promoter at the 5’ operon boundary, the presence of several additional promoters distributed through the operon raises the possibility that some genes are transcribed separately. Future analysis of the transcriptome of 95NR1 or other *E. coli* that carry homologs of M.Eco95NR1IX will be necessary to correctly determine transcriptional start-sites. Notably, SMRT sequencing also offers the potential for characterising complete polycistronic mRNAs using the Iso-Seq protocol (Sharon et al. 2013; Tilgner et al. 2014).

Characterising the genetic differences between strains is highly important for determining the evolutionary history of bacterial populations, tracking clinical outbreaks and identifying functional mutations which contribute to virulence and antibiotic resistance (Holt et al. 2008; Loman and Pallen 2008; Petty et al. 2014; Bartley et al. 2016). Currently, Illumina is the platform of choice for studying single nucleotide variation, due to its capacity for accurate high throughput sequencing of hundreds or thousands of strains. In this study the SNP profile between 95JB1 and 95NR1 was identical using PacBio or Illumina data, whereas only PacBio could accurately resolve the mobile genetic element content of both strains. Of particular importance to our understanding of STEC biology was the complete resolution of two tandemly arrayed Stx2 prophages encoded by 95NR1 and not 95JB1. The complete sequence of three full-length Stx2 prophages was determined in 95NR1 highlighting the difficulty of resolving multiple similar prophage genomes within a draft assembly. Although a highly accurate WGS method for subtyping *stx* genes has been developed (Ashton et al. 2015), the reliance of this method on short read sequencing data means it lacks the discriminatory power to determine whether multiple copies of the same subtype are due to multiple insertions by different Stx-converting phage or as a result of gene duplication. Our study highlights how population genomic studies of STEC outbreaks or global collections could benefit from SMRT sequencing and/or bioinformatics approaches that take into account mobile genetic element heterogeneity.

Although the ability of SMRT sequencing to resolve large mobile genetic elements is currently unparalleled, it is important to recognise that small plasmids are easily lost during the library preparation for PacBio. On the PacBio instruments the size distribution of the SMRTBell sequencing libraries influences read length performance. For example, short DNA library fragments preferentially load into the sequencing wells (Zero-mode wave guides) on the SMRT Cell limiting the long read potential of the sequencing library. In order to maximise the lengths of reads prior to sequencing it is necessary to filter small fragments from the library and narrow its size distribution. In this study targeted DNA size selection was achieved using the BluePippin instrument (http://www.sagescience.com). Using BluePippin small DNA fragments are separated from larger fragments enabling the collection of sequencing libraries with narrower size distributions. Both 95NR1 and 95JB1 were sequenced using BluePippin size-selected 20 kb SMRTBell libraries. As small plasmid DNA was visible in the original DNA extraction (data not shown) it appears that DNA fragments representing the missing colicin plasmids of 95NR1 and 95JB1 were filtered from the sequencing library during targeted size selection.

Here we have analysed the complete genome sequences of two clonal STEC O111:H-strains from a historical outbreak using PacBio sequencing. In addition to resolving complex prophage regions to highlight the precise genetic differences between these strains, we also identified more than 4000 differentially methylated chromosomal sites implicating a MTase that was active in 95NR1 but not 95JB1. We linked the differences in methylome to the mobilization of an IS*1203*-like insertion sequence in 95JB1, highlighting the importance of characterising IS differences when comparing two closely related bacterial strains. These results reveal that in addition to acquiring a small number of SNPs and losing two tandemly arranged Stx2 prophages, 95JB1 has also lost the activity of a novel MTase that may influence the transcription of several hundred genes. PacBio SMRT sequencing has enormous potential to reveal the genetic and epigenetic heterogeneity within a clonal population. Further genomic analysis of IS and prophages within closely related STEC strains will further build our understanding of short-term evolution within the context of an outbreak.

## Methods

### Genome Sequencing and assembly

Genomic DNA (gDNA) from the *Escherichia coli* strains 95NR1 and 95JB1 was sequenced on a PacBio RSII instrument (University of Queensland Centre for Clinical Genomics; UQCCG) using two SMRT cells, a 20 kb insert library and the P6 polymerase and C4 sequencing chemistry. *De novo* assembly of the raw PacBio sequencing data was done using the hierarchical genome assembly process (HGAP version 2) and Quiver (Chin et al. 2013) from the SMRT Analysis software suite (version 2.3.0 – http://www.pacb.com/devnet/) with default parameters. Following *de novo* assembly all contigs were visually screened for overlapping sequences on their 5’ and 3’ ends using contiguity (https://github.com/mjsull/Contiguity)(Sullivan et al. 2015). Overlapping ends, a characteristic feature of the HGAP assembly process, were manually trimmed based on sequence similarity and the contigs were circularised. Circularised contigs (Chromosome and plasmids) were then subjected to a polishing phase, were the raw PacBio sequencing reads were mapped back onto the assembled circular contigs (BLASR (Chaisson and Tesler 2012) and quiver) to validate the assembly and resolve any remaining errors. Following multiple rounds of polishing an additional improvement step was required to resolve single nucleotide insertion and deletions errors associated homopolyer tracts. Reads from the publically available Illumina sequence data for 95NR1 (SRA accession: SRR953500) and 95JB1 (SRA accession: SRR954273) were aligned to their respective genomes using bwa version: 0.7.12-r1039(Li and Durbin 2009) and a corrected consensus was called using Pilon version 1.18 (Walker et al. 2014). A total of 42 indels were corrected in 95NR1 and 30 indels in 95JB1. 95NR1 was further processed as the incorrect distribution of reads between two tandemly, integrated Stx prophages initially resulted in a misassembly characterised by a contig break. This misassembly was manually corrected, and verified by re-aligning the raw reads to the complete chromosome.

### Methylome analysis

The detection of methylated bases and clustering of modified sites to identify methylation-associated motifs was performed using the RS_Modification_and_Motif_analysis.1 tool from the SMRT analysis package version 2.3.0. Raw reads were aligned to the complete genomes of 95JB1 and 95NR1 and interpulse duration (IPD) ratios were calculated using PacBio’s *in silico* kinetic reference computational model (http://www.pacb.com/wp-content/uploads/2015/09/WP_Detecting_DNA_Base_Modifications_Using_SMRT_Sequencing.pdf).

To compare the CTGCAG motif distribution of Phi9 and Phi12 with the rest of the chromosome, the sequence for each strand was split into 1000 bp segments with a 250 bp overlap using Bedtools v2.17.0 (Quinlan and Hall 2010). Analysis of the mean distribution of CTGCAG motifs per segment within these genomic regions was performed as previously described using a custom analysis of variance (ANOVA) R script, which adjusts for heteroscedasticity (Forde et al. 2015).

### SNP analysis of 95NR1 and 95JB1

To determine the number and position of unique SNPs that differentiate 95JB1 and 95NR1, Illumina reads were simulated from the complete genomes of 95NR1 and 95JB1 and aligned to the genome of *E. coli* O111:H-strain 11128. SNP calling and Indel prediction was performed using Nesoni and the Nesoni n-way pairwise comparison method was used to identify SNPs conserved in all three strains (http://www.vicbioinformatics.com/software.nesoni.shtml). Additionally, a reference free SNP analysis was performed by direct comparison of the complete genome of 95JB1 and 95NR1 using MUMmer version 3.2.3 (Kurtz et al. 2004).

### Genome annotation and comparative genomics

Initial gene calling and automated functional annotation of 95NR1 and 95JB1 were performed using Prokka (Prokka: Prokaryotic Genome Annotation System – http://vicbioinformatics.com/) using a custom *Escherichia* genus database consisting of protein sequences from the EcoCyc website (http://ecocyc.org/). Putative phage encoding loci and IS elements were identified using PHAST (Zhou et al. 2011) and ISfinder (Siguier et al. 2006) respectively. Artemis Comparison Tool (ACT) (Carver et al. 2005), EasyFig (Sullivan et al. 2011) and Circos (Krzywinski et al. 2009) were used to visually compare the genomes and methylomes of 95NR1 and 95JB1 and identify regions of similarity and difference. Methytransferase genes and restriction modification systems were identified and annotated by comparison of all coding sequences from 95JB1 and 95NR1 against the REBASE database (Roberts et al. 2010). Promoter sequences were predicted using BPROM(Li 2011).

### The functional characterisation of CRARCAG methylation

*In silico* prediction of promoter sequences was done using Neural Network Promoter Prediction version 2.2(Reese 2001) and KEGG gene ontologies (KOs) were assigned using BlastKOALA (Kanehisa et al. 2016). The proximity of the CRARCAG motifs to all protein coding regions was determined using custom python scripts. Protein coding regions that did not contain a methylated CRARCAG motif within 300 bp of its start codon were excluded from the analysis. For the remaining protein coding regions, 300 bp of sequence upstream of their respective start codons was screened for putative promoter regions. CRARCAG motifs located within predicated promotor regions where identified, and KOs were assigned using BlastKOALA.

## Acknowledgements

This work was supported by Discovery Grant DP170103962 from the Australian Research Council. J.C.P. is a National Health and Medical Research (NHMRC) Senior Principal Research Fellow; S.A.B. is a NHMRC Career Development Fellow.

